# TB-annotator: a scalable web application that allows *in-depth* analysis of very large sets of publicly available *Mycobacterium tuberculosis* complex genomes

**DOI:** 10.1101/2023.06.12.526393

**Authors:** Gaetan Senelle, Christophe Guyeux, Guislaine Refrégier, Christophe Sola

## Abstract

Tuberculosis continues to be one of the most threatening bacterial diseases in the world. However, we currently have more than 160,000 Short Read Archives (SRAs) of *Mycobacterium tuberculosis* complex. Such a large amount of data should help to the understanding and the fight against this bacterium. To accomplish this, it would be necessary to thoroughly and comprehensively examine this significant mass of data. This is what TB-Annotator proposes to do, combining a database containing all the diversity of these 160,000 SRAs (at least, SRAs with a reasonable read size and quality), and a fully featured analysis platform to explore and query such a large amount of data. The objective of this article is to present this platform centered on the key notion of exclusivity, to show its numerous capacities (detection of single nucleotide variants, insertion sequences, deletion regions, spoligotyping, etc.) and its general functioning. We will compare TB-Annotator to existing tools for the study of tuberculosis, and show that its objectives are original and have no equivalent at present. The database on which it is based will be presented, with the numerous advanced search queries and screening capacities it offers, and the interest and originality of its phylogenetic tree navigation interface will be detailed. We will end this article with examples of the achievements made possible by the TB-Annotator, followed by avenues for future improvement.

## 1 Introduction

Tuberculosis remains the most common cause of death from a single infectious bacterium. The World Health Organization’s End TB Strategy targets to reduce TB deaths by 95% and to cut new cases by 90% between 2015 and 2035 [1]. An estimated 6.4 million tuberculosis (TB) cases and 1.6 million deaths have occurred in 2021, compared to 5.8 million TB cases and 1.5 million deaths in 2020 worldwide: the eradication of this disease does not seem to be within reach. Tuberculosis has been decimating humanity since antiquity [2]. Its infectious agent was identified by Robert Koch in 1882, and it was later called the *Mycobacterium tuberculosis* bacillus. Since the discovery of Calmette and Guérin’s biliary vaccine in the early 20th century and the one of antibiotics shortly thereafter, one of the most promising advances has undoubtedly been the complete sequencing of the bacterium’s genome in 1998 [3]. Since then, its sequence has been constantly studied, which has allowed to lift the veil on the complexity of its evolution.

Thanks to genomic studies, it has been shown that tuberculosis in humans is mainly caused by the members of the *Mycobacterium tuberculosis* complex (MTBC). *Mycobacterium tuberculosis sensu stricto* includes 5 lineages: L1 (Indo-Oceanic or EAI), L2 (East-Asian or Beijing), L3 (East-African-Indian or CAS), L4 (Euro-American), and L7 (Ethiopia). *Mycobacterium africanum* includes two other lineages: L5 (West African 1) and L6 (West African 2) [4]. More recently, two new lineages, namely the L8 [5] restricted to the African Great Lakes region and the L9 [6], have been described. Genomic studies have also allowed significant progress to be made in terms of the mechanisms at the origin of its virulence, as well as those explaining the increasing resistance to antibiotics [7, 8].

The cost of sequencing has also dropped dramatically over the past 25 years, making the acquisition of new sequences common and affordable. Whole Genome Sequencing (WGS) technologies also referred to as Next Generation Sequencing data (NGS) have made it possible to collect more than 160,000 MTBC genome sequences publicly available on the NCBI SRA database [9], the well known repository for high-throughput sequencing data. Such database provides an essential source of information for the study of the evolution of tuberculosis, but tools are obviously lacking to fully exploit it. The objective of our new platform is not to offer *yet another pipeline method* that does more or less what is already being done but do propose a new paradigm in systems biology of tuberculosis. Its justification comes from the following facts.

First of all, as detailed in Section 3.1, existing tools mainly focus on specific subjects like resistance prediction and genotyping of bacterial isolates. Their aim is not to provide a global and complete approach based on all available SRAs, but something specific on data provided by the user. For instance, some pipelines have been developed to predict potential drug-resistance, like the acknowledged and widely used TB-profiler and/or Phyresse [10,11], Indeed, such pipelines do not have tools to explore and analyse genomic characteristics at a large scale. They usually focus on one type of markers, see Sect. 3.1. They may be poorly equipped with global analytic tools. Some pipelines use a whole bunch of tools, difficult to maintain, difficult to run, and does not allow to scale to the whole MTBC genomes.

As will be seen in Section 3.1, a number of tools are not, by construction and reason to be, specific to *M. tuberculosis*. They therefore gain in generality what they lose in specificity. If they can be applied in a non-specific way to a variety of bacteria, they conversely cannot take advantage of decades of specific knowledge accumulated about the genomic characteristics of the MTBC genomes. In a more subtle way, each pipeline produces different results in incompatible format which makes the construction of a global analysis pipeline, integrating several complementary tools, rather complicated to achieve. In the same vein, it is frequently difficult to compare private strains with already publicly available genomes. To sum up, to set up fully integrated workflows from scratch, that allow an accurate and meaningful analysis of NGS data from clinical MTBC strains, still requires programming expertise, and trained bioinformatics staff. This constrains the application of MTBC NGS analysis to specialized laboratories, leads to a large diversity of analysis pipelines with group-specific solutions that seriously complicates standardized comparison of results.

This explains why we developed the *TB-Annotator* database, that contains informations on raw phylogenetical markers and annotations. It implements information retrieval methods used for text datasets, which have been proven useful for large-scale corpora. This database is at the same massive scale as the SRA one, and it integrates most of known genomic characteristics of MTBC genomes (lineage markers definitions, regions of difference, insertion sequences…). As such, it is the largest pre-analyzed database specific to the *Mycobacterium tuberculosis* complex. Moreover, this database comes with an ergonomic and original interface, allowing to make complex and advanced queries or to create complex filters, and to investigate with a yet unachieved depth, any specific phylogenetic tree of your choice. As will be seen below, TB-annotator has already allowed us to revisit a certain amount of knowledge on two lineages of tuberculosis [12, 13]. While access is currently restricted (interested readers can apply to the authors for access), it is intended to become public soon.

To the best of our knowledge, such a platform that would process massive amount of data is not yet available for *Mycobacterium tuberculosis* complex genomes. Concretely, this large amount of still unexploited spatio-temporal genomic diversity data could constitute in a near future a gold-mine of a yet hidden knowledge, that would allow to perform a kind of archeological dive into the past of specific infectious diseases history. With more than 160,000 SRA available, and constantly updated, the field is quite mature, and part of the work will consist in being able to discriminate what are the information that will really improve our global understanding of the tuberculosis pandemia, from the data that will be either useless or contribute to the well-known overlearning process. By doing so, one of the final aim of this evolving project could be to design for tuberculosis a similar project than the one that was launched just before the Covid-19 sanitary crisis, whose ambition was to perform real-time tracking of pathogens [14], or even to more ambitiously create a platform that would allow to perform deep-learning for an improved characterization, diagnostic or treatment of TB diseases applied to patient or communities, in relation to genomic data analysis.

The rest of this article is structured as follows. In the following section, we present TB-Annotator in detail, both in terms of the database and its interface. We discuss its originality in Section 3, showing that its objectives are different from the tools currently used by the community. Examples of its implementation are also given in this section, for illustration purposes. This article ends with a perspective, in which the interest of the TB-annotator is recalled, with new studies that we wish to carry out or are currently in progress.

## 2 TB-annotator description

TB-Annotator is made up of two main components. First the pipeline which massively analyse all MTBC publicly available genome. And second, the analysis platform to explore and search this data.

### 2.1 Pipeline architecture

First of all, the pipeline uses Snakemake [15] as the execution engine. Snakemake is designed to build reproductible and scalable data pipeline by leveraging a declarative workflow definition and modern execution platforms such as cloud (for example see Kubernetes [16]). This workflow engine was selected over cloud native workflow engines such as Argo Workflows [17] for its ease of use, ease of deployment on personal desktops, and the facilitated integration with common bioinformatics tools.

The pipeline starts with a list of accession numbers provided by the user. Sequences are then downloaded from Sequence Read Archive (SRA) [18], with fastq-dump, as paired-end or single-end FASTQ files. This first step of the pipeline is throttled to reduce the number of simultaneous connections to the NCBI. FASTQ files contain sequencing reads with the sequence and a quality score for each base, the PHRED score [19]. These scores are used in the reads pre-processing and mapping steps to assess the reads quality, therefore FASTQ is the only supported input format.

For non-publicly available sequences, the pipeline accepts compressed FASTQ files. As a convention, sequences from private sources are given a custom identification number starting with the prefix *CUS* (for example CUS000001). The pipeline is preconfigured to detect this naming scheme, and to analyse *CUS* sequences at the same time as SRA sequences.

The pipeline DAG (directed acyclic graph) and computations are organized to maximize data reuse and incremental enhancements. It allows to perform new analyses and computations without doing again costly steps like SRA download.

### 2.2 Read preparation

Genome coverage is estimated based on the number of reads in the FASTQ file. To ensure the speed of the pipeline remains constant, the FASTQ file is randomly downsampled to a configurable estimated genome coverage using SeqKit [20].

Fastp [21], an all-in-one and high performance tool, is used for quality control on the reads and to pre-process them. In the pre-processing part, the adapters used in library preparation process during sequencing are trimmed. Low quality bases at ends of reads are cut and reads with too many low quality bases are removed. For paired-end reads that are overlapping, they are corrected based on quality score if they are not a perfect reverse-complement. Some advanced corrections are also performed like polyG trimming and UMI preprocessing [22].

Following the read quality control and pre-processing, a report is outputted in html and json formats. The report contains informations and statistics about the different parts of the reads filtered, base contents, and duplication rates. This report can be used to evaluate the sequencing process, and detect problems like contaminated reads, base content biases and over-represented sequences. Original FASTQ files are removed after this process to reduce disk space usage.

### 2.3 Read mapping

Processed reads are mapped on a reference genome (NC 000962.3 by default) using BWA-MEM [23]. BWA-MEM is one of the fastest mapper [24], while being close to Novoalign in terms of accuracy [25]. Duplicate reads are marked using SAMTOOLS [26] for the next tools in the pipeline, including freebayes, to prevent bias in the variant calling due to PCR duplicates that might cause the algorithm to identify an error during the amplification as a true variant [27].

The sequence alignment is saved in CRAM format, a compressed format based on the reference sequence [28]. The resulting file is small enough to be kept on hard drive at low cost, and to be able to perform further analysis or to execute an updated pipeline in the future without executing the download and pre-processing step a second time. This optimization is made possible thanks to Snakemake ability to continue partially executed workflows. The computing time is decreased by using the sequence alignment as a basis for all the subsequent steps of the pipeline, instead of executing analysis on raw reads.

Finally statistics are computed with SAMTOOLS using the sequence alignment file, like mean read depth and the proportion of covered bases. This information can be used to assess the quality of the sample and the quality of sequencing. This information will also be used in the rest of the pipeline for statistical analysis.

### 2.4 Variant calling and annotation

Snippy [29] is used for variant calling, which is one of the top performing pipeline for bacterial genomes [30]. Snippy takes as input the already aligned reads in compressed CRAM format instead of letting it aligns the read (by default with bwa-mem too) to bam format. Local realignment was not performed before variant calling because it had a low impact with the tools used [30].

Snippy starts with the use of Samclip [31] on aligned reads to avoid false SNP calling close to structural variations. It works by removing clipped reads inside contigs. The resulting alignment mainly contains long aligned reads and thus providing higher confidence in variations called. Variants are then called with Freebayes [32]. The final output of this step is a VCF file.

Some regions of the MTBC genomes are known to be difficult to map including mobile elements, Proline-Glutamate (PE)/Proline-Proline-Glutamate (PPE) genes and repetitive regions. These regions usually contain a number of soft-clipped reads that are removed by samclip, but some of them are kept, for example reads corresponding to insertions sequences mapped to IS regions of the MTBC reference. These problematic regions are removed in number of studies [33]. An additional filtered VCF file is outputted in which variants falling in these regions are removed.

Variants are annotated using SnpEff [34], a variant annotation and effect prediction tool. SnpEff annotates and predicts the effects of genetic variants (such as amino acid changes). We identify variants using their SPDI notation [35], this allows to have a globally unique identifier for each variant. The alternate representation in which the Deletion field is a string containing the literal sequence to delete [35] is used to avoid the need for reference sequence when reading the SPDI.

### 2.5 Read clipping analysis

When a read partially matches the reference sequence, the aligner usually removes the unmatched part, this process is called clipping. Two types of clipping exist: (1) hard clipping where the removed part is indeed removed from the resulting alignment file, and (2) soft clipping where the removed sequence is kept and its position is signaled in the CIGAR string. By default BWA-mem uses soft clipping for primary alignment. A large number of clipped reads at the same positions is usually a sign a of Structural Variant (SV), and it is used for general SV calling [36].

We call *clipping signal* the position where a configurable amount of reads is clipped. A clipping signal can be on the left side or the right side of reads (Figure 4). In the clipping signal detection step, a custom Python script creates the list of clipped reads, the position of the left clipping and right clipping and the clipped sequence. The list of clipping signals is created using *bedtools groupby* [37] on the output of the Python script.

**Fig 1.**
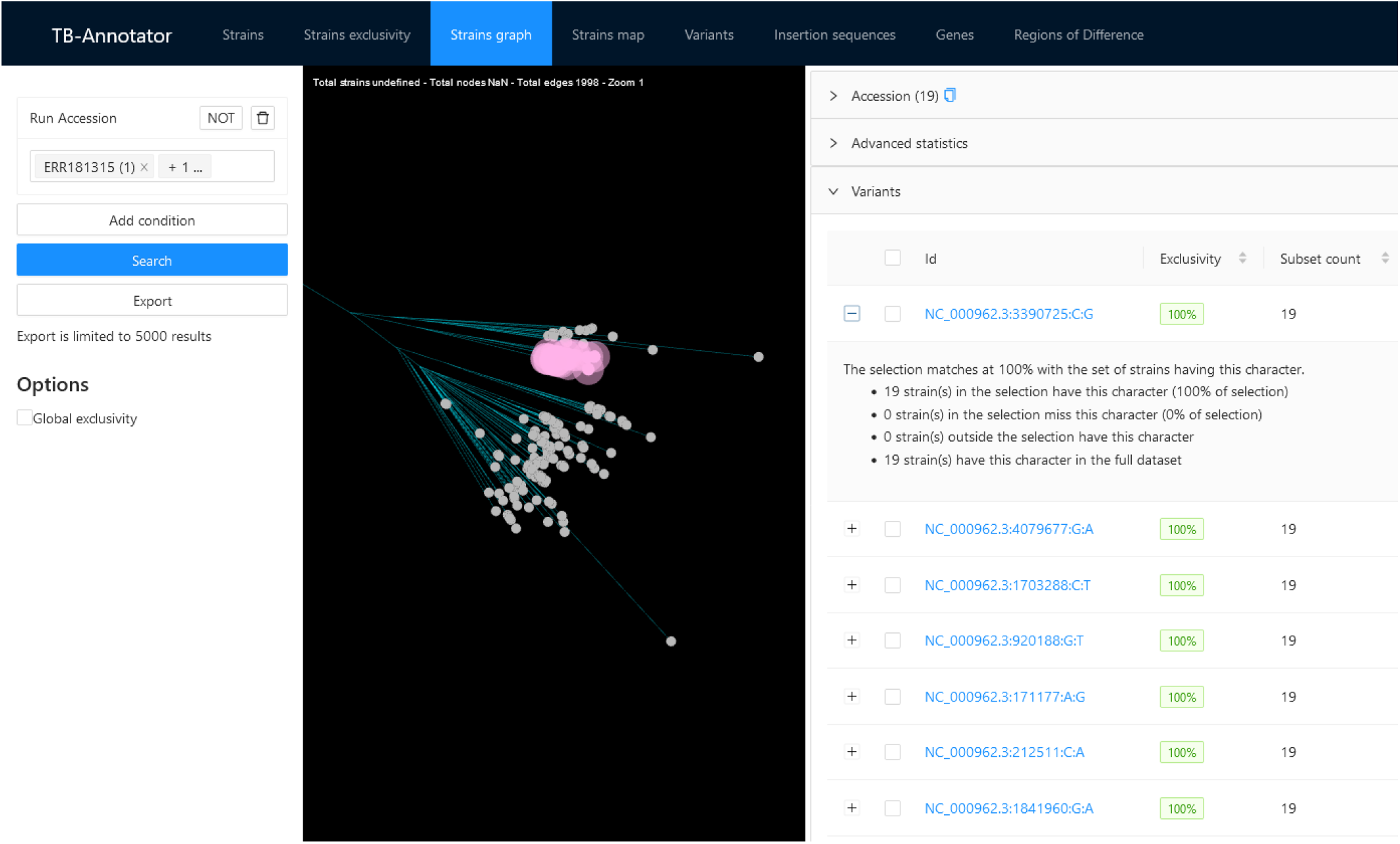
Graph visualization

**Fig 2.**
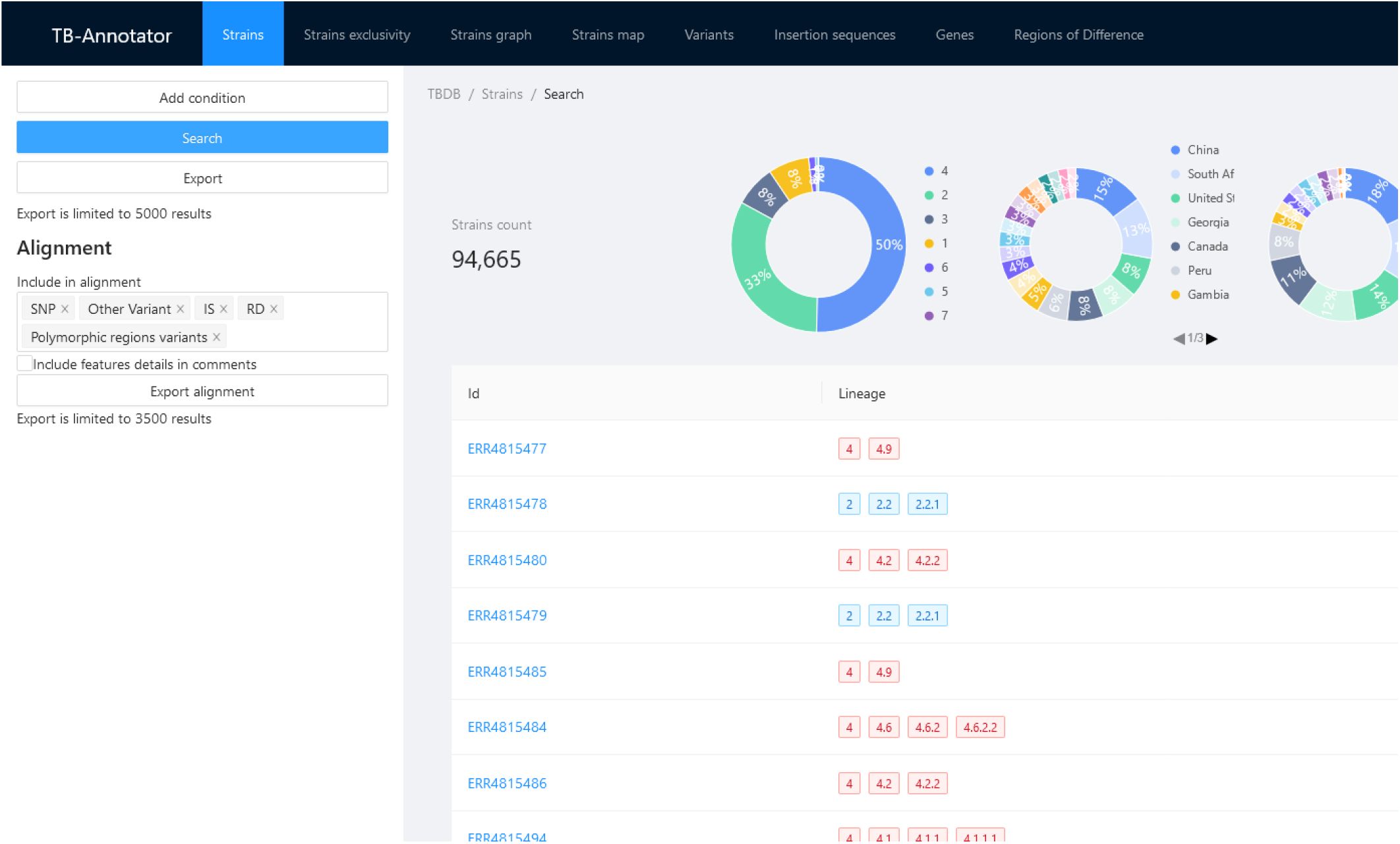
Database overview

**Fig 3.**
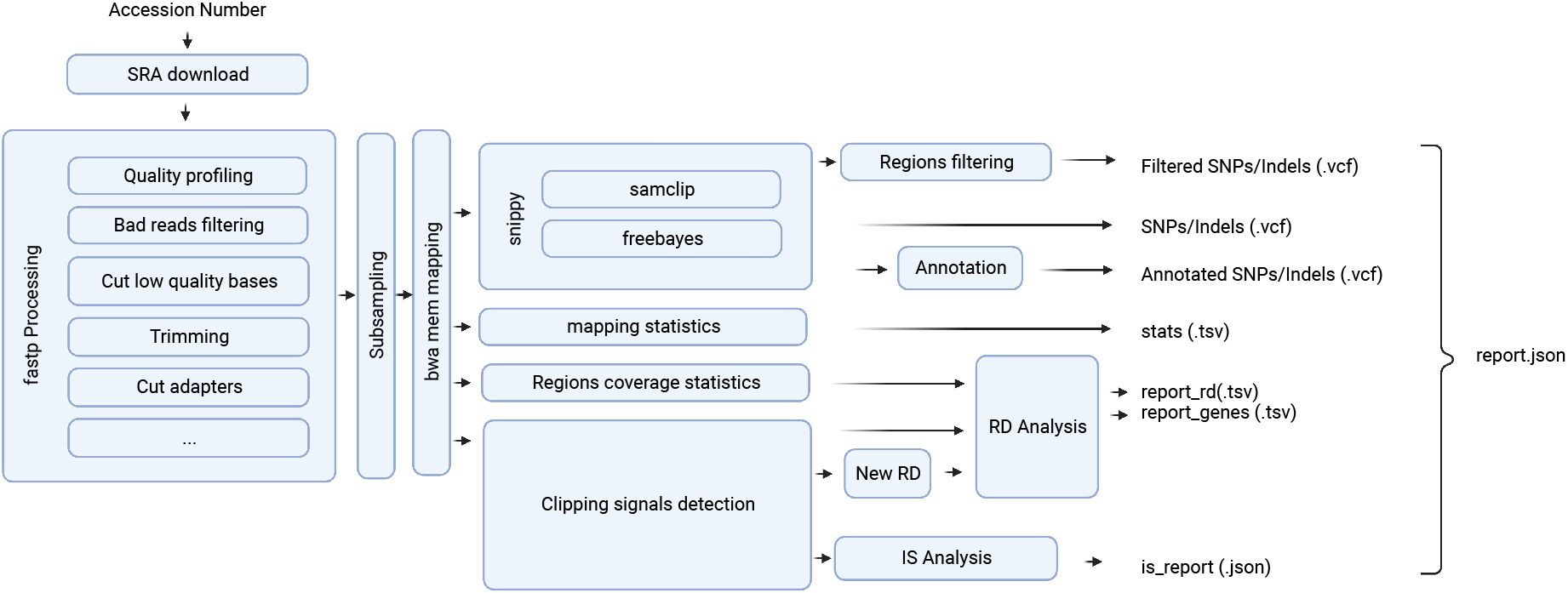
Pipeline overview

**Fig 4.**
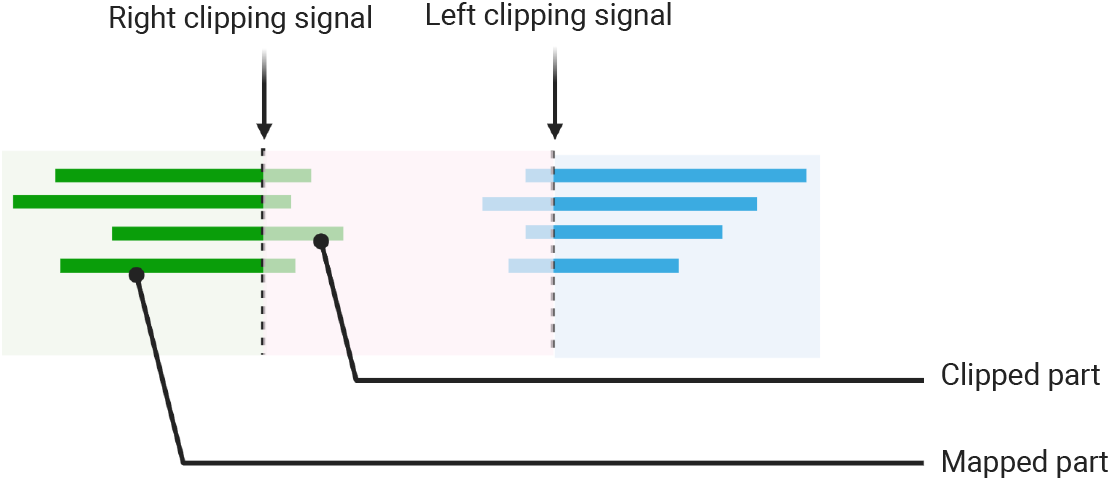
Clipping signals

This step is only a pre-processing for subsequent steps. The clipping information can be used to detect SV such as deletion and insertions. And contrary to the general case of SV detection, we can use this information to detect known MTBC genome features more easily and more precisely.

### 2.6 Detection of the present/absence of known RD and genes

Known Regions of Difference (RD) were manually collected from more than 20 studies, curated and checked. The positions were converted to positions on the reference genome H37Rv NC 000962.3. When this position was not available in the original article, it was inferred using various methods like mapping the PCR junction, mapping example strains containing this deletion, or extracting the position from other references like *M*.*bovis*.

This step focuses on assessing the presence/absence of genes and large RD (RD much smaller than read length, like pks 1/15 7bp insertion, are found by the variant calling step [38]). A read depth histogram is computed on the aligned reads using BEDTOOLS [37]. For each region (RD and genes) multiple statistics are computed like mean, median, min, max coverage and the ratio 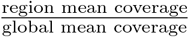 Most of existing approaches for *in silico* RD detection uses these metrics [6, 39, 40]. With this method, we miss various information, for example: if the region is fully deleted or not, if the region is part of a larger deletion or precisely deleted, if it is probable that the noticed drop in read depth is indeed related to a deletion. To overcome some of these issues, we compute the percentage of very low read depth in the given region using a configurable threshold. The resulting value can be used as an indication of partially deleted gene for example.

The second method is based on clipping signals extracted from a previous step. We use these signals to find what we call *high quality deletions*, which are regions for which we are highly confident that they are precise deletions (and not false positives due to lower read depth in the region, for example). Clipping signals close to the sides of the region are searched using a configurable flank. Clipped part is aligned on the opposite side of the region as described in Figure 5. When a configurable number of sequences precisely aligns on both sides, we mark the region as *high quality*.

**Fig 5.**
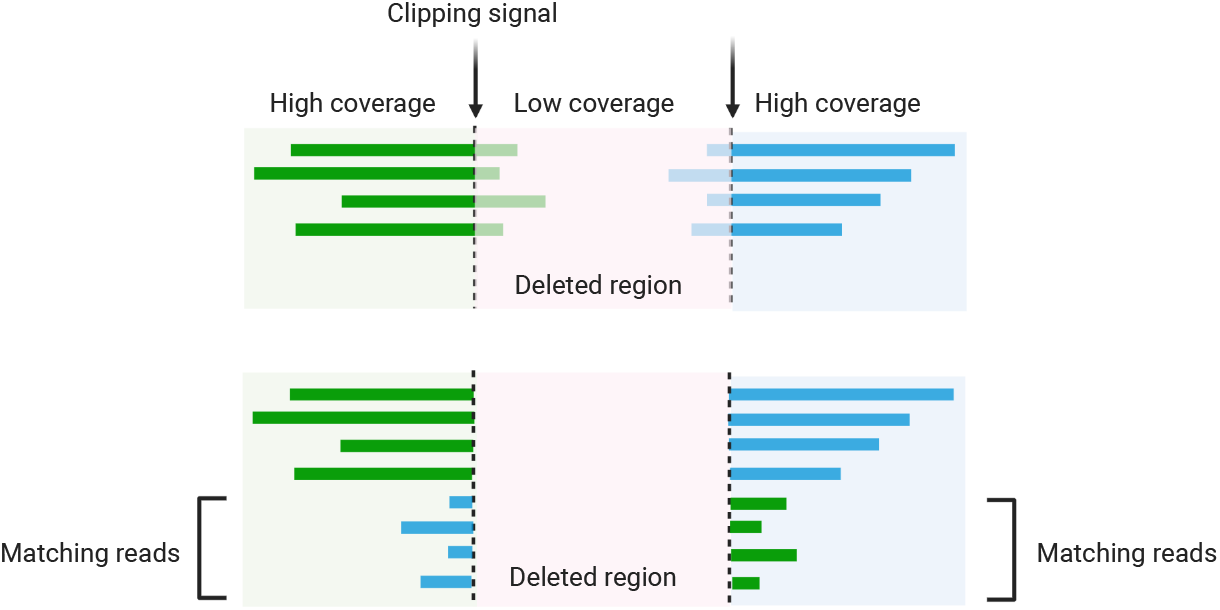
Mapping of clipped sequences

Flanks are used when searching clipping signals mainly due to the way reads are aligned. The clipping signal is not always close to the end of deletions. This is the case for example when the sequence before the start of the deletion equals the sequence at the end of the regions. This case is illustrated on Figure 6. Because we know approximately where the two clipping signals are for a given region, this decreases the risk of mapping clipped sequences at a wrong position in the genome. Not all regions can be considered high quality, like RD falling in larger deletions. These regions are still detected using read depth variation like explained earlier. This case is illustrated in Figure 7.

**Fig 6.**
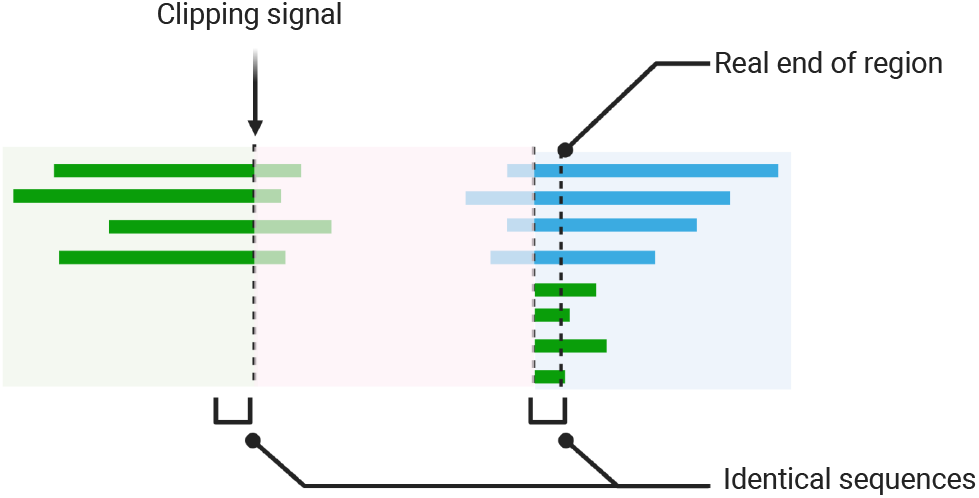
Difference between clipping signal and ends of deleted regions

**Fig 7.**
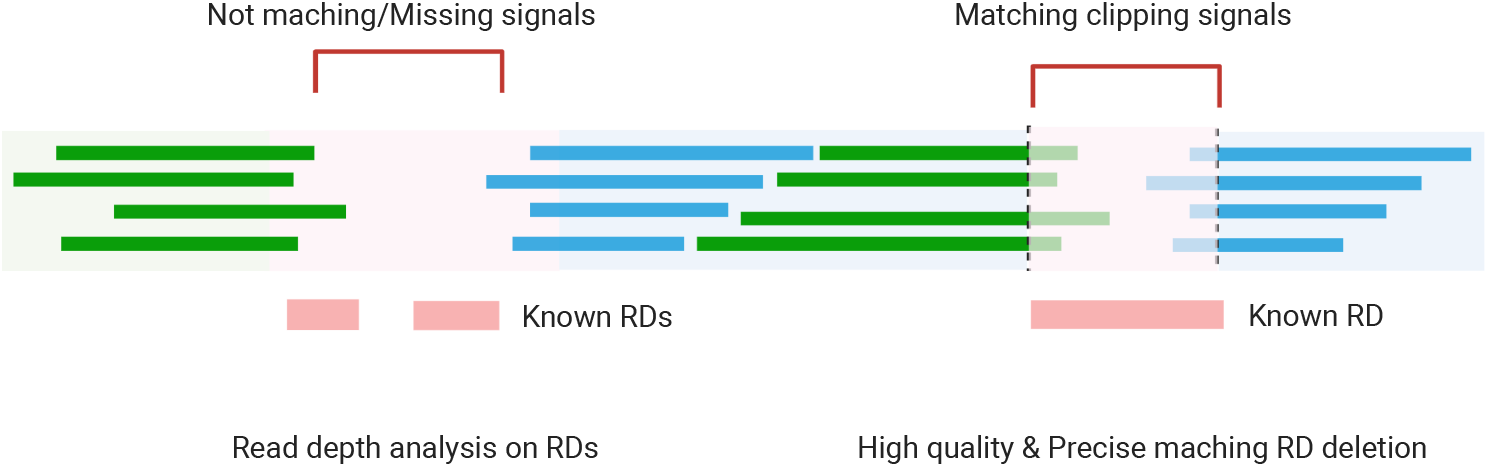
Different methods of detection of regions deletions

The pipeline comes with two list of regions, genes and known RDs, while additional regions are configurable with bed files. For each list of regions, a TSV report is created, with all statistics described earlier and a score indicating if the region is considered as *high quality*. At the end of this step, all the searched regions are outputted. The final choice (if a region is deleted or not) is left to the last step by the creation of the final report. This allows to keep detailed information on regions for further analysis.

### 2.7 Detection of new RD

The pipeline allows the detection of RDs that are not documented in existing studies. To ensure the consistency and precision of positions detected across all strains analysed, this step is also based on previously detected clipping signals. Possible regions of deletion are searched by creating pairs of consecutive clipping signals, and a name is generated for each of these pairs. These regions are then analysed at the same time as known regions to increase pipeline speed.

### 2.8 Detection of CRISPR

Reads are mapped using BWA-MEM [23] on the sequences of all known spacers. For each spacer, instead of outputting binary presence of the spacer, we output statistics to help evaluate the probability of the presence or absence of the spacer.

### 2.9 Detection of insertion sequences

We collected sequences as fasta files from 22 known MTBC insertion sequences (https://wwwis.biotoul.fr/).

Clipping signals detected in a previous step are used to detect insertion sequences. Pairs of clipping signals separated by a configurable small distance are searched. For each pair of clipping signals, clipped sequences from the reads are extracted (Figure 8) and mapped using BWA-MEM [23] on a fasta file containing all known insertion sequences.

**Fig 8.**
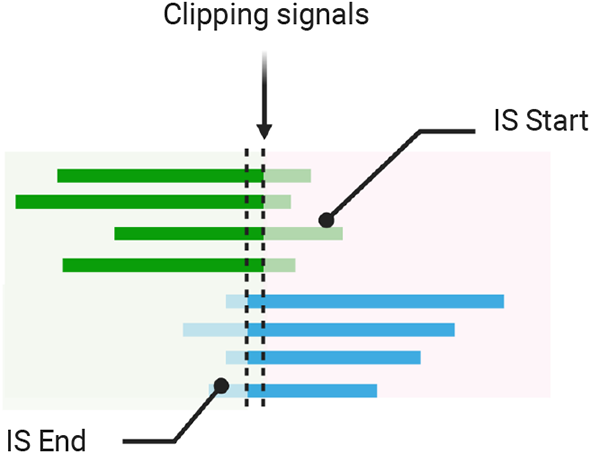
Insertion sequence detection using clipping signals

The matching insertion sequence is determined by analysing the resulting mapping file. We find the first sequence where (1) clipped sequences are mapped on both sides of the reference insertion sequence, (2) left clipped sequences align on the same position, and (3) right clipped sequences align on the same position. This process ensures that the start and the end of the insertion is coherent. This method also allows to detect IS insertion orientation depending on which side of the reference insertion sequence reads are mapped. When the inserted sequence is a subsequence of an IS, we also store the start and the end of the subsequence relatively to the reference IS.

For insertion sequences contained in the reference, RD detection algorithm is used to determine if the insertion sequence is deleted. This detection is made possible thanks to the clipping signal algorithm. Achieving this detection would not be feasible using RD algorithms relying on read depth, as reads from other insertion sequences may map to the deleted region.

### 2.10 Final report

The output of all the steps of the pipeline are gathered in a single JSON report containing:

- the list of variants found and their annotations,
- the list of missing genes,
- the list of missing region of differences,
- coverage statistics for each gene and RD,
- the list of insertion sequences,
- and the quality report from Fastp.

### 2.11 Pipeline execution

We run the pipeline with 128 threads on a Dell PowerEdge R740 hosted at the Mésocentre de calcul de Franche-Comté. The system is using a 16 core Intel(R) Xeon(R) Gold 6226R CPU @ 2.90GHz with 128GiB of RAM.

MTBC SRA are analysed by batches of 15,000, each batch taking approximately one week to complete. Task scheduling is handled by Snakemake using the greedy scheduler to decrease the job graph computation time.

### 2.12 Data search

In order to search the analysed strains, all the report data is enriched and indexed in an inverted index. Inverted index was chosen due to the fact that most queries consist of searching strains having a certain character (like a specific variant), and such queries are very efficient using inverted indexes.

An indexing pipeline, developed in C# using Dataflow API, is used to read the genomic pipeline JSON reports, and to index the data in Elasticsearch 8. The report is first read in memory, splitted in 4 types of data (strain, variant, RD, IS), annotated and finally indexed in separate indices. The indexing performance is close to 20 reports per second using a 8 cores AMD Ryzen 7 2700 Processor @ 3.20GHz with 32GiB of RAM, with a local instance of the database.

### 2.13 Feature exclusivity computation

The feature exclusivity allows to select a subset of clinical isolates (called foreground set) among the set (called background set) of all genome sequences publicly available on the NCBI SRA database, and to determine if the features of those isolates (variants, RD, IS) are exclusive to the subset. TB-Annotator is able to run such analysis on very large amount of strains by leveraging the inverted index global ordinals. Using a custom script, the Jaccard coefficient is computed to be used as a score on every feature of the strains.

### 2.14 External annotations during indexing

The Snakemake pipeline is only based on genomic data, all additional enrichments are done during the indexing.

Each variant indexed in the database is annotated for resistance based on TBDB [41] and WHO [42] data. When a SNP is a known lineage marker, the SNP is annotated with a reference to the author [43–48]

For each isolate, the indexer starts by collecting all public metadata available in the NCBI database. Results are cached for faster re-indexation and to limit number of request to NCBI API. The isolate is then annotated for resistance based on variants annotation associated to this isolate.

Genes, for their parts, are annotated using NCBI NC 000962.3 metadata.

### 2.15 Web platform

TB-Annotator Web platform allows to browse, search and run advanced analyses on the database from the user browser. User is able to perform fast queries on most attributes of strains, variants, insertion sequences, genes and RD using the inverted index.

The creation of alignments to be used in tools like RAxML is done by querying the database in batches, and dynamically computing a map between each document and the features it contains. The map is then used to generate an alignment in FASTA format. In case where other features than SNP are exported, a binary alignment is used. The details of columns is also exported at the end of the FASTA file in comments.

For the *strain* graph feature, user phylogenetic trees in Newick are first parsed in the browser. The tree is then rendered using WebGL and equal-angle method layout algorithm also implemented in the browser. The name of each node in the graph is then extracted to be sent to the server to be used as list of accession. It allows the use of feature exclusivity computation against arbitrary phylogenetic trees. The list of all displayed strains is used as a background set for exclusivity comparison (the whole database can also be used as background set).

## 3 Discussion

The aim of this section is to show that, on the one hand, the TB-annotator is original and complementary to existing tools and, on the other hand, that it is useful, having already enabled us to make new discoveries within the MTBC.

### 3.1 Comparison with existing tools for *M.tuberculosis* genome analysis

In this section we will compare TB-Annotator to existing web platforms and other databases available.

#### 3.1.1 WGS data analysis tools

TBprofiler [49] is another next-generation sequencing analysis tool for *Mycobacterium tuberculosis* that focuses on detecting drug resistance mutations. TBprofiler also provides a web interface, but its scope is limited to drug resistance analysis. TB-Annotator goes beyond this by allowing analysis and visualization of insertions, difference regions, and missing genes, thus providing a more comprehensive analysis of the genomes.

CASTB (Comprehensive Analysis Server for the Mycobacterium tuberculosis Complex [50]) is a database and web server that allows the analysis of whole-genome sequences of *M. tuberculosis* complex strains. The tool provides information on drug resistance and lineage identification through a series of analyses, such as SNP-based phylogeny and prediction of drug resistance mutations. However, unlike TB-Annotator, CASTB is not designed to be used for real-time analysis of large-scale genomic data, and it does not have a comprehensive annotation pipeline to detect genomic variations beyond known resistance mutations.

Mykrobe [51] is a suite of tools for the identification of drug resistance in bacterial pathogens, including *M. tuberculosis*. The tool provides predictions for resistance to first-line and second-line drugs, and it also allows the identification of lineage for isolates. However, mykrobe is not designed for comprehensive annotation of genomic variations beyond resistance mutations.

Compared to these tools, TB-Annotator provides a comprehensive pipeline for the detection of genomic variations, including SNPs, insertions, deletions, and large-scale genomic rearrangements. Additionally, TB-Annotator allows for the annotation of genomic variations beyond known drug resistance mutations, enabling researchers to study the evolution and transmission of *M. tuberculosis* strains in greater detail. Furthermore, TB-Annotator provides a user-friendly web platform that allows for real-time analysis and visualization of large-scale genomic data. Overall, TB-Annotator stands out as a unique tool that provides a comprehensive and user-friendly approach to the annotation and analysis of genomic data in the context of *M. tuberculosis* research.

#### 3.1.2 Genetic profiling

MIRU-VNTRplus [52] is an online analysis tool designed for the identification, typing, and phylogenetic comparison of MTBC strains. The tool is based on the typing method called MIRU-VNTR (Mycobacterial Interspersed Repetitive Units-Variable Number Tandem Repeats), which is a PCR-based molecular typing technique. MIRU-VNTRplus provides a platform for comparing new isolates to a reference database, allowing researchers and clinicians to better understand the genetic diversity, epidemiology, and transmission patterns of tuberculosis. This tool is therefore very different from ours: it focuses on a single subject of analysis (MIRU-VNTRs), of exclusively experimental origin, and it handles only one isolate at a time (the one submitted by the user). Our tool does not have as input experimental MIRU-VNTR extraction data, but read files. It is unable to analyse these MIRU-VNTRs, as the size of the reads makes it impossible to count copies of motifs of interest. However, it does produce all possible analyses on sequencing data, and provides a comparative analysis of all available genomes (with sufficient quality and read size).

SITVITWEB [53] is an online database and analysis tool for Spoligotyping-based genotyping of *Mycobacterium tuberculosis* complex (MTBC) strains. Spoligotyping is a PCR-based method that detects the presence or absence of specific spacer sequences in the direct repeat region of the MTBC genome. SITVITWEB allows users to query the database with new Spoligotype patterns, providing information on global distribution, phylogenetic classification, and epidemiological context. The tool aids researchers and public health professionals in understanding the population structure, geographical distribution, and transmission patterns of tuberculosis strains.

#### 3.1.3 Gene annotations and expression profiles

TBDB [54] is an integrated database containing information about *M. tuberculosis* genes, proteins, and drug resistance mutations. It serves as a valuable resource for researchers by consolidating data from various sources and providing tools for data analysis and visualization. Although TBDB offers a rich collection of genomic information, it does not offer an analysis pipeline for whole-genome sequencing data, as TB-Annotator does. TB-Annotator not only analyzes and annotates genomic data but also provides advanced search capabilities and a platform for running advanced analyses on the resulting data.

TubercuList [55] and MycoBrowser [56] are online databases containing annotated genomic information for *Mycobacterium tuberculosis*. These resources allow for gene and protein exploration, providing detailed information on gene function, gene ontologies, and metabolic pathways. However, they do not provide the analysis, visualization, and search capabilities that are central to TB-Annotator. TB-Annotator’s comprehensive approach to whole-genome sequencing data analysis, along with its advanced search and visualization features, sets it apart from these databases.

PATRIC [57] is an extensive online resource for bacterial pathogens, offering tools for genomic and comparative analysis. PATRIC covers a wide range of bacterial species, including *M. tuberculosis*. While it provides valuable data and analysis tools for researchers working on various bacterial pathogens, it is not specifically tailored for *M. tuberculosis* and does not provide the same depth of analysis, visualization, and strain-specific data as TB-Annotator. TB-Annotator is designed exclusively for *M. tuberculosis*, enabling a more focused and comprehensive analysis of this particular organism.

BioCyc [58] is a collection of pathway and genome databases that cover various organisms, including *Mycobacterium tuberculosis*. BioCyc offers valuable information on metabolic pathways, gene function, and genome annotations. However, it does not provide the comprehensive genome analysis, visualization, and search capabilities that TB-Annotator offers. TB-Annotator is designed specifically for *M. tuberculosis*, allowing it to provide a more detailed and focused view of the genomic features of this organism, along with advanced search and analysis options.

#### 3.1.4 Other partly comparable websites

Enterobase [59] is a publicly accessible database of bacterial pathogens, including *Mycobacterium tuberculosis*. It allows users to submit sequencing data and compare it with other isolates to identify outbreak clusters and track the spread of antimicrobial resistance. While it has some similar functionality to TB-Annotator, it does not provide the same level of annotation and analysis of genomic features. Additionally, Enterobase does not currently include all available *M. tuberculosis* sequencing data, whereas TB-Annotator incorporates a large portion of the publicly available SRA data.

PathogenSeq [60] is a web-based platform that provides tools for the analysis of microbial genomic data, including *M. tuberculosis*. It allows users to upload their own data or use pre-loaded datasets and provides tools for assembly, annotation, and comparative genomics. However, unlike TB-Annotator, PathogenSeq is focused more on providing raw data analysis tools rather than pre-annotated data. Additionally, it does not include some of the features of TB-Annotator, such as the ability to search for specific genomic features, investigate phylogenetic trees, and perform feature exclusivity analysis.

Nextstrain [14] is a web-based platform that provides real-time tracking of global infectious disease outbreaks, including M. tuberculosis. It allows users to view phylogenetic trees of viral and bacterial sequences, track the spread of outbreaks, and identify new genetic variants. While it provides some similar functionality to TB-Annotator, it does not provide the same level of annotation and analysis of genomic features, nor does it include some of the search and filtering options of TB-Annotator. Additionally, Nextstrain is focused more on tracking the spread of disease in real-time, whereas TB-Annotator is more focused on providing a comprehensive database of M. tuberculosis genomic features.

CNGBdb [61], finally, is a publicly accessible database of genomic data, including *M. tuberculosis*. It provides tools for searching and analyzing genomic data, as well as data visualization and comparison. While it has some similar functionality to TB-Annotator, it does not provide the same level of annotation and analysis of genomic features, nor does it include some of the search and filtering options of TB-Annotator. Additionally, CNGBdb does not include all available *M. tuberculosis* sequencing data, whereas TB-Annotator does.

#### 3.1.5 Summary

In conclusion, TB-Annotator is a unique tool that stands out for its ability to provide a comprehensive analysis of *M. tuberculosis* genomes. Unlike most of the existing tools, TB-Annotator integrates several steps that range from quality filtering of reads to annotation and indexing of detected variants, insertion sequences, and regions of differences. Furthermore, TB-Annotator has a web-based interface that allows easy visualization and mining of data from the database. Compared to other tools, TB-Annotator offers several additional features, such as the detection of new RDs, the ability to detect insertion sequences, and the computation of feature exclusivity. Moreover, TB-Annotator provides external annotations during indexing that improve the resistance annotation of indexed variants. Finally, TB-Annotator is capable of analyzing a large number of genomes simultaneously, making it ideal for population-based studies.

Overall, TB-Annotator offers a powerful and user-friendly platform for exploring M. tuberculosis genomic diversity, and its unique features make it a valuable addition to the existing tools. By providing a comprehensive analysis of genomic data, TB-Annotator will facilitate better understanding of TB epidemiology and resistance patterns, which will ultimately help in the design of more effective control strategies. Additionally, its web-based interface and the ability to analyze large datasets will make it a useful resource for researchers and public health professionals alike.

### 3.2 Examples of using the TB-annotator to answer specific questions

#### 3.2.1 Connection between two historical tuberculosis outbreak sites in Japan, Honshu, by a new ancestral Mycobacterium tuberculosis L2 sublineage [62]

In this study, we describe a historically endemic ancestral sublineage of L2, known as AAnc5, based on samples collected in the Tochigi Prefecture in central Japan. This sublineage is closely related to the Japanese G3 group identified in 2012 and is presumed to be named L2.2.A in a recent review. The findings of this study strengthen the phylogenetic relevance of this sublineage within the global L2 evolutionary history, showing that it was historically transmitted in several Japanese cities.

Tuberculosis is an ancient disease in human history, but its exact emergence in Asia remains uncertain. Early TB outbreaks in Japan may be connected to population migrations between the 5th century BC and the 3rd century AD. Tuberculosis was known to be present in ancient Japan under the name of rôga, which was used in Chinese medicine. Traces of this disease can be found in ancient Chinese medical texts. A critical aspect of this study is the use of the TB-Annotator pipeline, which has greatly facilitated the analysis of large amounts of data contained in Sequence Read Archive (SRA) files. By analyzing over 50,000 characters, including repeated sequences and single nucleotide polymorphisms (SNPs), TB-Annotator enables a more in-depth examination of the complex historical phylodynamics of all *Mycobacterium tuberculosis* complex (MTBC) lineages. This tool has played a significant role in obtaining the results presented in this study and in mapping the endemic L2 ancestral sublineage from Japan onto the global MTBC phylogeny.

Among the specific characteristics of AAnc5, the authors describe a non-synonymous mutation in the rpoC gene. This mutation is generally found in epidemiologically successful isolates that also contain specific rpoB gene mutations. The use of TB-Annotator has been crucial in identifying such unique features within the AAnc5 sublineage.

Estimations of the coalescence of AAnc5 sublineage landmarks range from 280-310 years to 760-800 years before the present. The Tochigi Prefecture is famous for its Ashio copper mine, which began operations in the early 17th century. The Ashio mine could have been a location for AAnc5’s expansion and diversification.

In conclusion, the TB-Annotator pipeline has been instrumental in mapping an endemic L2 ancestral sublineage from Japan onto the global MTBC phylogeny and designating it as Asia Ancestral 5 (AAnc5). This sublineage possesses many specific characteristics that distinguish it from other ancestral sublineages described so far in L2. The discovery of AAnc5 and the extensive insights provided by TB-Annotator open new avenues for research into the history of L2 in Southeast Asia.

#### 3.2.2 Tuberculosis in Nigeria: new insights from whole genome sequencing data (submitted work)

In this study, we investigated the genomic diversity of *Mycobacterium tuberculosis* lineages in Nigeria, with a focus on *M. africanum* (L5-L6) distribution. TB-annotator, an essential tool in the analysis, provides a deeper understanding of the genomic diversity and the rarer sub-lineages. The study reveals a fully diversified L5 lineage, while the L6 lineage is poorly represented. The absence of certain sub-lineages raises questions and will require further clarification. The low presence of L5-L6 in Nigeria suggests a possible disappearance of *M. africanum* in the region, a topic that has been debated in recent years.

TB-annotator plays a crucial role in analyzing these lineages, as it helps identify new isolates and their distribution. For example, the study found that the number of known isolates for the 511212 sub-lineage doubled, allowing for a better understanding of its specificity. The tool also aids in understanding the presence of certain sub-lineages in Nigeria, such as one-third of the L5 isolates being found in the country. This information is valuable for further research and could potentially lead to the discovery of new trends and patterns in the distribution of tuberculosis strains.

The study also discusses the importance of monitoring drug-resistance in Nigeria. A decade after the initial work, researchers found that 39.49% of patients were resistant to at least both INH and RIF, leading to an MDR rate well above 40%. This highlights the need for ongoing surveillance and the implementation of effective tuberculosis control programs. TB-annotator proves to be essential in this regard, as it helps assess drugresistance status and determine the prevalence of resistant strains in specific regions.

In summary, this research investigates the genomic diversity of *Mycobacterium tuberculosis* lineages in Nigeria, emphasizing the significance of TB-annotator in obtaining a comprehensive understanding of these lineages and their drug-resistance profiles. The study highlights the potential disappearance of *M. africanum* in Nigeria and the increasing importance of drug-resistance surveillance in the country. TB-annotator serves as a valuable tool for researchers to uncover new insights, track the spread of specific sub-lineages, and ultimately contribute to the development of more effective tuberculosis control strategies.

### 3.3 Examples of using the TB-annotator to answer more general questions: the case of the L5

The submitted article entitled “An updated evolutionary history and taxonomy of *Mycobacterium tuberculosis* lineage 5, also called *M. africanum*” focuses on lineage 5 (L5) of the *Mycobacterium tuberculosis* complex, which is a significant cause of tuberculosis in West and Central Africa. Despite the importance of L5, it has been poorly represented in public databases, which has made it challenging to reconstruct its evolutionary history and taxonomy. The authors aimed to build an exhaustive collection of representative L5 genomes and analyze their structure using TB-Annotator.

The study provides a considerable increase in the phylogenetical analysis of L5, offering a better description of basal groups, the identification of new reliable SNPs at each subbranch, and an integrated phylogenetic reconstruction of the entire L5 lineage. The authors note that diversification, i.e., transmission, was pervasive in the beginning of L5 history but diminished in later evolutionary periods. This ancient diversity could still be underestimated due to under-sequencing of the geographic regions concerned by this lineage.

TB-Annotator proved to be useful in the phylogenetic analysis of L5 by providing a finer hierarchical classification and identification of new signature SNPs. The authors note that the phylogenetic resolution is more difficult towards the root, which is expected, but this can be solved with the so-called exclusive variants approach. A notable exception to the overall lack of L5 genomic dynamism is the insertion of IS6110 between spacers 30 and 35 in the CRISPR locus, which is a feature of the L5.1 sublineage.

The study still suffers from some limitations, including a lack of geographic origin information for many samples, absent or scarce data for many countries of Africa, and few genomic studies linking human genetics predisposition to bacterial genetics. However, the authors were able to provide a list of 49 SNPs that allow for the resolution of the phylogenetic tree of L5 with three main branches: L5.1, L5.2, and L5.3 (new). They also linked specific IS6110 insertion/deletion events with particular sublineage emergence, such as the loss of spacers 21-24 in relation to the loss of IS6110 in L5.1.

Finally, the authors discuss the historical timeframe and geographical area of L5 emergence in a wider evolutionary context, suggesting that L5 and L6’s highly geographically-constrained patterns may be related to the domestication and migration of *Bos taurus* and *Bos indicus*, respectively. The authors suggest that L4 may have emerged in relation to a North-african *Bos taurus* domestication, and L1-L5-L6 MTBC lineages could have been related to *Bos indicus* domestication and introduction into Africa via East-Africa and the Arabian peninsula. The study provides a more precise picture of some of the evolutionary steps of L5 diversification and offers insight into the intricate relationships between human and animal life-styles, from hunter-gatherer to more recently urban life-styles.

In conclusion, the study offers a comprehensive analysis of L5, providing new insights into its evolutionary history and taxonomy. We were able to use TB-Annotator to provide a finer hierarchical classification and identification of new signature SNPs, which is useful in further studies of tuberculosis caused by *M. africanum*. Although the study still suffers from some limitations, it highlights the need for further genomic studies in underrepresented regions to better understand the virulence evolution of all MTBC lineages.

## 4 Conclusion

As such, the app is already mature. It incorporates 102,001 strains out of the 167,062 available on NCBI, at the time of this writing. The strains not integrated either have too short reads (the majority) or do not pass our quality filters. In detail, we find a majority of lineage 4 (50%), followed by L2 (33%), L3 and L1 (8% each), the other lineages being marginal (1,122 strains of L6, 452 of L5, 96 of L7, 2 of L8 and 12 of L9). It also contains animal strains, namely *orygis, pinipedii, microtii, dassie* (*Procavia capensis*), *caprae, bovis* (including BCG), *mungi* and *chimpanzee*.

The web application contains two main functionalities. The database part allows multi-criteria searches for the presence of traits of interest: SNP (2,209,467 variants), presence of IS, gene or RD, as well as searches by country, by bioproject, by resistance… The alignment corresponding to the search for these traits can be extracted, and if a tree is built from it by an external application, the newick file can be integrated into the interface in order to exploit the second main functionality, namely visualization and data mining within this tree. By clicking on a strain (SRA), one can visualize all the information about it contained in the database. More interesting, we can make multiple selections (of clades) and see the characters in common and exclusive. We can make multi-criteria searches of characters and visualize in the tree the strains that have them. Finally, we can navigate in the tree (zoom, translation). Because of the various optimization layers mentioned above, all these operations are done in a fluid and almost instantaneous way.

The application is still under intense development. Among the new functionalities being integrated, we can mention the production of an estimate of the spoligotype per SRA (the spacer box is darker the more likely it is to be in the CRISPR locus), the production of a distance matrix per pair of strains in the context of multiple selections, as well as the assembly of reads into contigs. The latter is not currently used, but it should eventually allow the detection of genes that are not present in the H37Rv reference.

For the future, the main objective is to first extend access to the platform to a larger number of private users, and then to move to a web application accessible to all. Such a deployment implies ensuring the scaling up, the security and stability of the platform, as well as its sustainability. Ideally, everyone should be able to deposit their own genomes, with all the guarantees that this implies. Finally, we would like to increase the number of references by not restricting ourselves to H37Rv, integrate long reads (PacBio, etc.), and why not extend the database to other bacteria.

*All computations have been performed on the Mésocentre de Franche-Comté super-computer facilities*.

